# Experimental landscape connectivity decreases temporal variability in communities over 24 years of assembly

**DOI:** 10.64898/2026.06.16.732628

**Authors:** Katherine A. Hulting, Lars A. Brudvig, Melissa A. Burt, Christopher R. Warneke, Ellen I. Damschen, Nick M. Haddad

**Affiliations:** W.K. Kellogg Biological Station, Michigan State University, Hickory Corners, Michigan, USA; Department of Integrative Biology and Program in Ecology, Evolution, and Behavior, Michigan State University, East Lansing, Michigan, USA; Department of Plant Biology and Program in Ecology, Evolution, and Behavior, Michigan State University, East Lansing, Michigan, USA; Division of Math and Science, Tusculum University, Greeneville, Tennessee, USA; Department of Integrative Biology, University of Wisconsin-Madison, Madison, Wisconsin, USA

**Keywords:** Connectivity, community assembly, variability, dispersal, beta diversity

## Abstract

Landscape connectivity is a key regulator of dispersal, which is an important process in community assembly. Theory predicts that connectivity may influence spatial and temporal patterns of community assembly; however, empirically evaluating the role of connectivity is nearly impossible due to the need to isolate its influence over long time frames and large spatial extents. We overcome these challenges through a large-scale, long-term connectivity experiment to test how connectivity affects plant community turnover and directionality of change over 24 years of assembly. Plant communities within connected patches had lower temporal variability in composition compared to plant communities within unconnected patches. Differences in composition between patches and the directionality of compositional changes were driven more by the amount of edge habitat in a patch and the time since the start of assembly. All community responses to connectivity were stronger for species with wind or unassisted dispersal compared to those with seeds dispersed by animals. Connectivity’s role in regulating local community dynamics is critical for understanding community assembly and increasingly relevant in an era of anthropogenic land-use change.

**Significance Statement:** Connectivity between habitat patches facilitates dispersal to localities, yet the impact of connectivity on local species assemblages is exceptionally challenging to isolate from other spatial changes over time. In a 24-year experiment, we found that connectivity stabilized local community composition as a higher number of species persisted across years within patches connected by corridors. Independent of connectivity, edge effects were more important for driving compositional differences between patches. Importantly, these patterns would not have been captured with short-term data or without controlling for confounding spatial changes. Our findings have broad conservation relevance. Anthropogenic landscape changes that result in a loss of connectivity or increased edge effects may disrupt local community assembly over time.

## Introduction

A central goal of ecology is to understand patterns in community assembly (1–4). Dispersal is a key process regulating community assembly, yet experiments testing dispersal’s role tend to directly manipulate dispersal by moving individuals or seeds (5–7), without testing the mediating role of landscape connectivity in facilitating dispersal (8). However, this focus limits progress in understanding assembly, as local communities exist in the broader context of regional landscape connectivity, which can make dispersal to local communities non-random (4, 9, 10). Integrating connectivity into our understanding of community assembly links local communities with regional dynamics, but, as of yet, few studies have experimentally tested the role of connectivity in assembly over timescales long enough to capture these dynamics.

Connectivity, or the extent to which a landscape facilitates movement (11), drives changes in local community assembly through increasing dispersal rates that influence the number of individuals and species arriving to a community from a regional species pool (10). The theory of island biogeography and metacommunity theory guide expectations for the consequences of dispersal rates on species richness in local communities (12, 13). Both theories predict that a higher dispersal rate due to connectivity increases colonization and reduces extinction, leading to a higher local (alpha) diversity in connected compared to isolated communities. These effects on colonization and extinction rates can be particularly evident for species with limited dispersal abilities, which may not be able to disperse to and persist in isolated communities (14–16). In addition, connectivity between local communities may also influence community assembly by maintaining species coexistence in local communities. High rates of dispersal allow species to persist in patches where they otherwise may not (mass effects) (17, 18), as well as alter biotic interactions that may determine coexistence patterns among species (10).

Despite the well-developed theory for the role of connectivity in community assembly, empirically evaluating community assembly across broad scales is nearly impossible due to the increased spatial extent required and the multiple landscape changes that co-occur with connectivity, such as increased habitat edge and decreased habitat amount (19). Additionally, because patterns in community assembly can change at different spatial scales or exhibit a lagged response to environmental fluctuations (20–22), small-scale or short-term studies may not be sufficient to capture trends in assembly, or may yield conclusions that are erroneous at larger scales (23). As a result, previous tests of connectivity on community assembly have been either unable to separate connectivity from other landscape changes, or unable to separate out these effects over long time periods (8). Interpreting the consequences of altered landscape connectivity for communities undergoing assembly requires long-term, landscape-scale experiments that disentangle these effects.

The role of connectivity for assembly dynamics over time can by evaluated through a framework of community convergence and divergence. Classical community assembly predicts that communities with similar environmental conditions assemble deterministically towards a stable state (1, 24), but other work emphasizes the potential for divergent community states (25, 26). These outcomes may be determined by differing rates of dispersal resulting in divergent rates of local colonization and extinction (27, 28). However, convergence or divergence between communities is not necessarily a steady trajectory towards an equilibrium state (29, 30). Communities may diverge in composition because of factors such as stochasticity in compositional change or variability in rates of colonization and extinction (30, 31). A loss of connectivity may impede species sorting and rescue effects (9, 18), which may increase temporal compositional turnover due to local extinctions. In contrast, increased dispersal linking communities may decrease temporal turnover within a local community by enabling species to persist in a patch (32). Capturing these temporal community dynamics requires long-term time series data (22) and is unable to be replaced by commonly used space-for-time substitution practices (33, 34).

These expectations for the consequences of connectivity on community assembly may also depend on the dispersal ability of species in a community and the spatial and temporal scale of the organisms and landscapes being considered (35). For plant communities, species with relatively long-distance dispersal abilities are more likely to disperse to isolated areas and, therefore, respond less to connectivity. In contrast, species with limited dispersal may show particularly strong responses to connectivity (35, 36). For plants, there is evidence that dispersal distances are greatest for animal-dispersed seeds, followed by wind- and gravity-dispersed seeds, which have the shortest dispersal distances (14, 37). Since landscape factors affect dispersal abilities in distinct ways, consideration of dispersal traits may provide insight into how communities may respond to habitat connectivity (15, 16, 35).

We overcome the challenges of spatial and temporal sampling by using large-scale experimental habitat corridors to test connectivity effects on 24 years of plant community assembly in longleaf pine savanna patches. Our experiment is composed of three types of habitat patches, all with equivalent habitat amount: patches connected by a corridor (‘connected’), unconnected patches with a high amount of edge (‘winged’), and unconnected patches with a low amount of edge (‘rectangular’) (Figure 1). This experimental design is ideal for addressing our research questions because we manipulate connectivity at a scale relevant to plant community dynamics, while also disentangling connectivity from other aspects of landscape change such as habitat amount, matrix quality, and edge effects (38, 39). Previous research in our experiment has found that connectivity increased colonization rates and decreased extinction rates, leading to higher plant species richness in connected than isolated patches (40). Using an extension of this dataset, we now ask new questions about community assembly over time:

**Figure 1.**
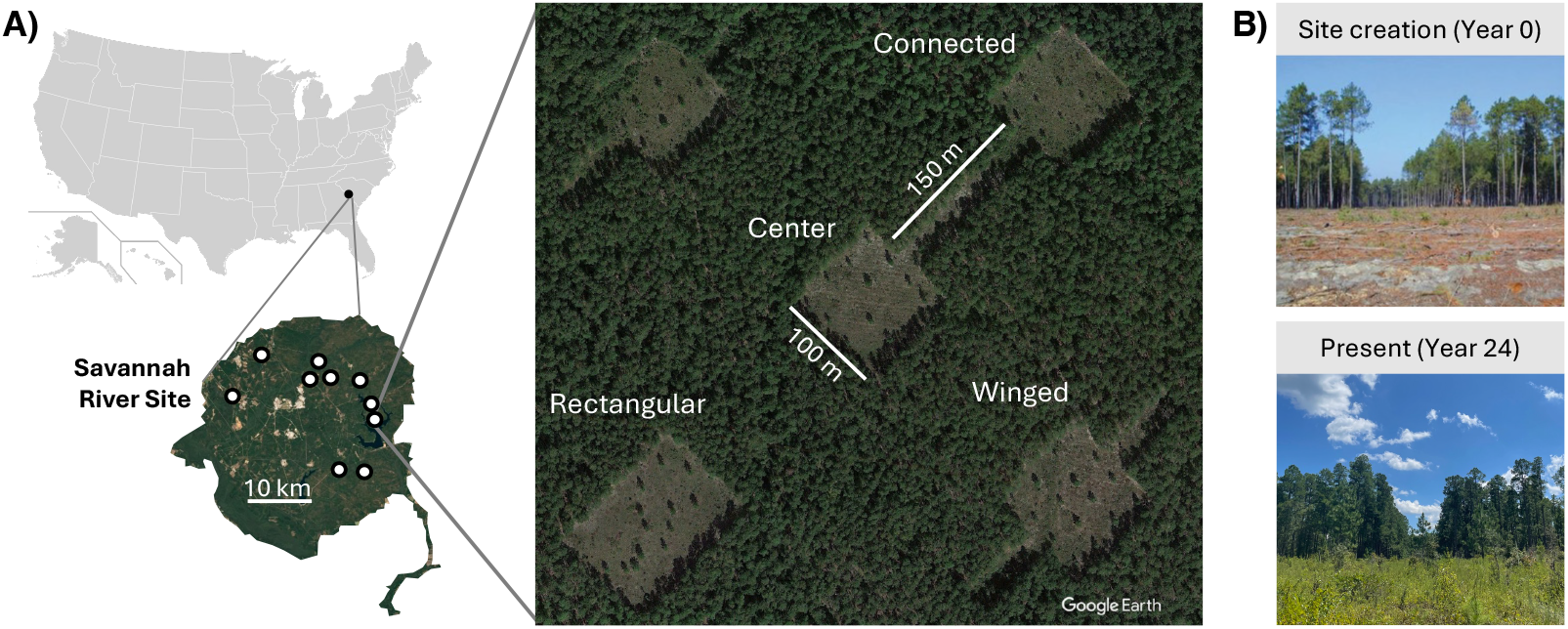
Design of experimental habitat corridors. A) Location of the 10 blocks within the Savannah River Site, South Carolina and aerial image of one block (Credit: Landsat/Copernicus & Google Earth 2022). Longleaf pine savanna patches were embedded in a pine plantation matrix and either connected to the center patch by a corridor (connected patch), unconnected with a high amount of edge habitat (winged patch), or unconnected with a low amount of edge habitat (rectangular patch). B) Example of longleaf pine savanna patch at the initiation of the experiment and 24 years into assembly (Credit: N.M. Haddad [top], K.A. Hulting [bottom]).

1. How does connectivity affect temporal dynamics within local communities, such as interannual variability in composition and assembly directionality?
2. Does connectivity or other landscape features cause local communities to diverge towards different community states compared to local isolated communities?
3. For each question above, does plant dispersal ability (i.e., dispersal mode) mediate the direction or timing of community assembly responses to connectivity?

## Results

### Temporal Variability in Composition Within Patches

Interannual variability in composition, or the degree to which communities have changed between consecutive years, was lower in connected patches compared to other patch types. To measure interannual variability in community composition, we calculated the degree of change in composition between years within each habitat patch using trajectory analysis (i.e., temporal turnover). By the midpoint of the time series, trajectory distance between consecutive years was significantly lower in connected patches compared to unconnected rectangular patches and unconnected winged patches (Figure 2A; Table S1-S3). In all patch types, trajectory distance between consecutive years decreased over time (about a 16.7% decrease over two decades of succession), indicating a reduction in year-to-year variability in community composition as communities assemble (95% CI: 6.7%-29.1% decrease) (Figure 2A; Table S2).

**Figure 2.**
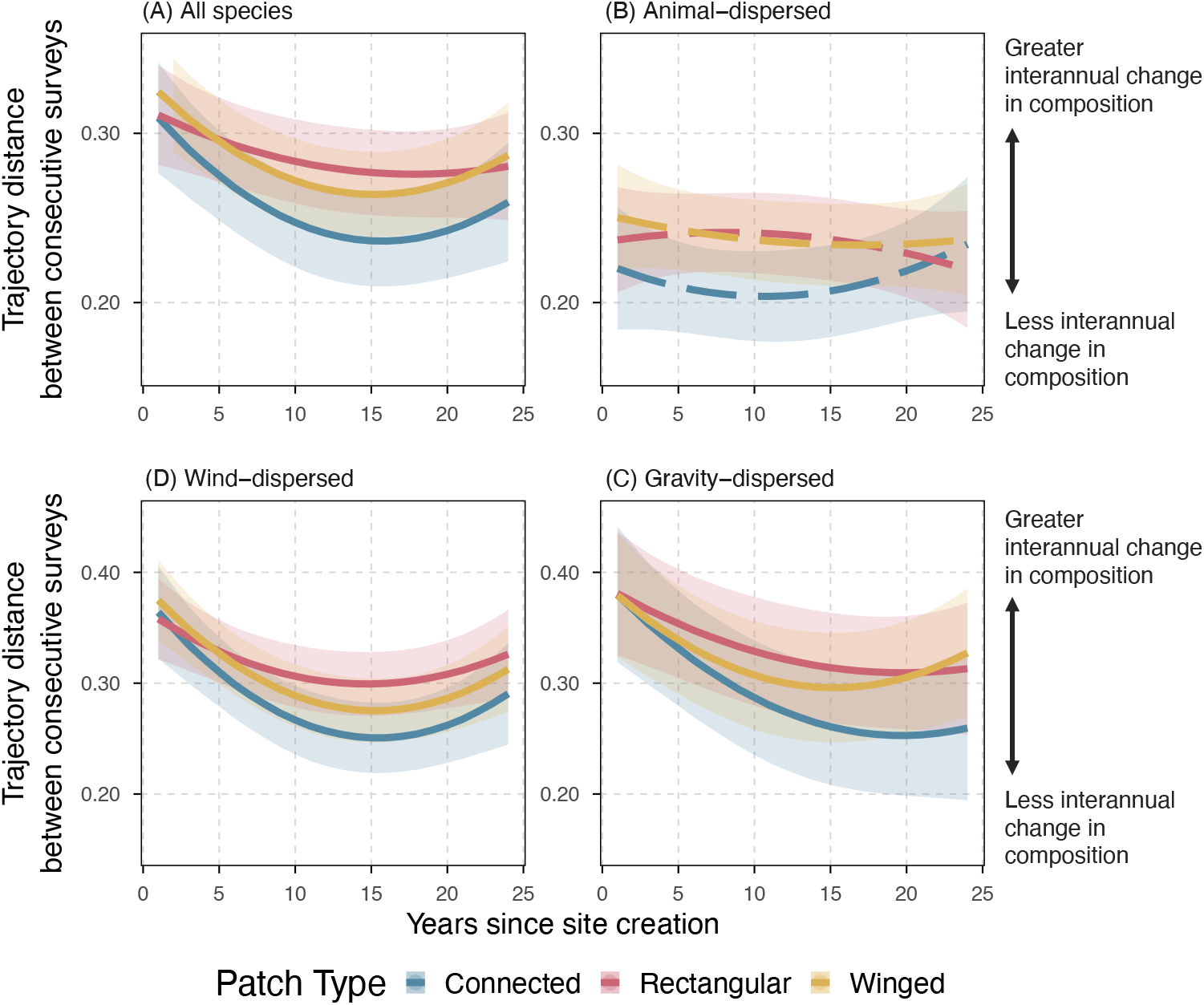
Temporal variability in composition within a patch: Community trajectory length (interannual change in composition) of (A) the entire plant community, (B) animal-dispersed species, (C) wind-dispersed species, and (D) gravity-dispersed species between consecutive surveys. The response variable is the distance of community change between t and t-1 sampling points. Higher values indicate more change in community composition between consecutive surveys and lower values indicate less change in community composition between consecutive surveys. Solid lines represent model predictions and shaded areas represent 95% CI. The dashed lines in (B) represent that the null model was the best fit model for animal-dispersed communities (Table S4). Note that the y-axis scales differ among panels.

A reduction in temporal compositional change within a community could be due to a fewer species gains and losses between years, or due to a higher number of species persisting between years, buffering changes. When partitioning compositional change into these components, we found that connected patches had a greater number of species persisting between consecutive years compared to other patch types (12% more species persisting compared to winged patches [95% CI: 2.9%-22.3%], 22% more species persisting compared to rectangular patches [95% CI: 12%-33%]). In contrast, the number of gains and losses between years was similar among patch types (Figure S1; Table S4-S6).

Among the three seed dispersal modes in our experiment (animal-dispersed, wind-dispersed, gravity-dispersed), communities of animal-dispersed species differed in temporal variability patterns from communities of wind and gravity-dispersed species. When subsetting the plant community to consist solely of wind-dispersed species or gravity-dispersed species, temporal compositional change was lower in connected patches compared to rectangular patches by the midpoint of the time series. In contrast, for communities of animal-dispersed species, patch type did not explain variation in temporal compositional change (Figure 2B; Table S1). Across time, compositional change between consecutive years decreased in communities of wind and gravity-dispersed species, but not in communities of animal-dispersed species (Figure 2B-D; Table S2, S3).

### Directionality in Compositional Changes

Across all patch types, community trajectories exhibited more directional change in composition at the beginning of assembly and more random changes in composition later in assembly (Figure 3A; Table S7, S9). We measured these patterns within patches using a metric of trajectory directionality to quantify whether temporal changes in composition in a multivariate space are more directional, linear paths or random, wandering paths, which could indicate how deterministically the community is assembling. Across all patch types, community trajectories became about 10% less directional in the second half of the time series compared to the first half (95% CI: 13.3%-6.0% less linear) (Figure 3A; Table S7). However, these temporal patterns in trajectory directionality were not impacted by connectivity or edge amount at any point in assembly (Figure 3A; Table S7, S8).

**Figure 3.**
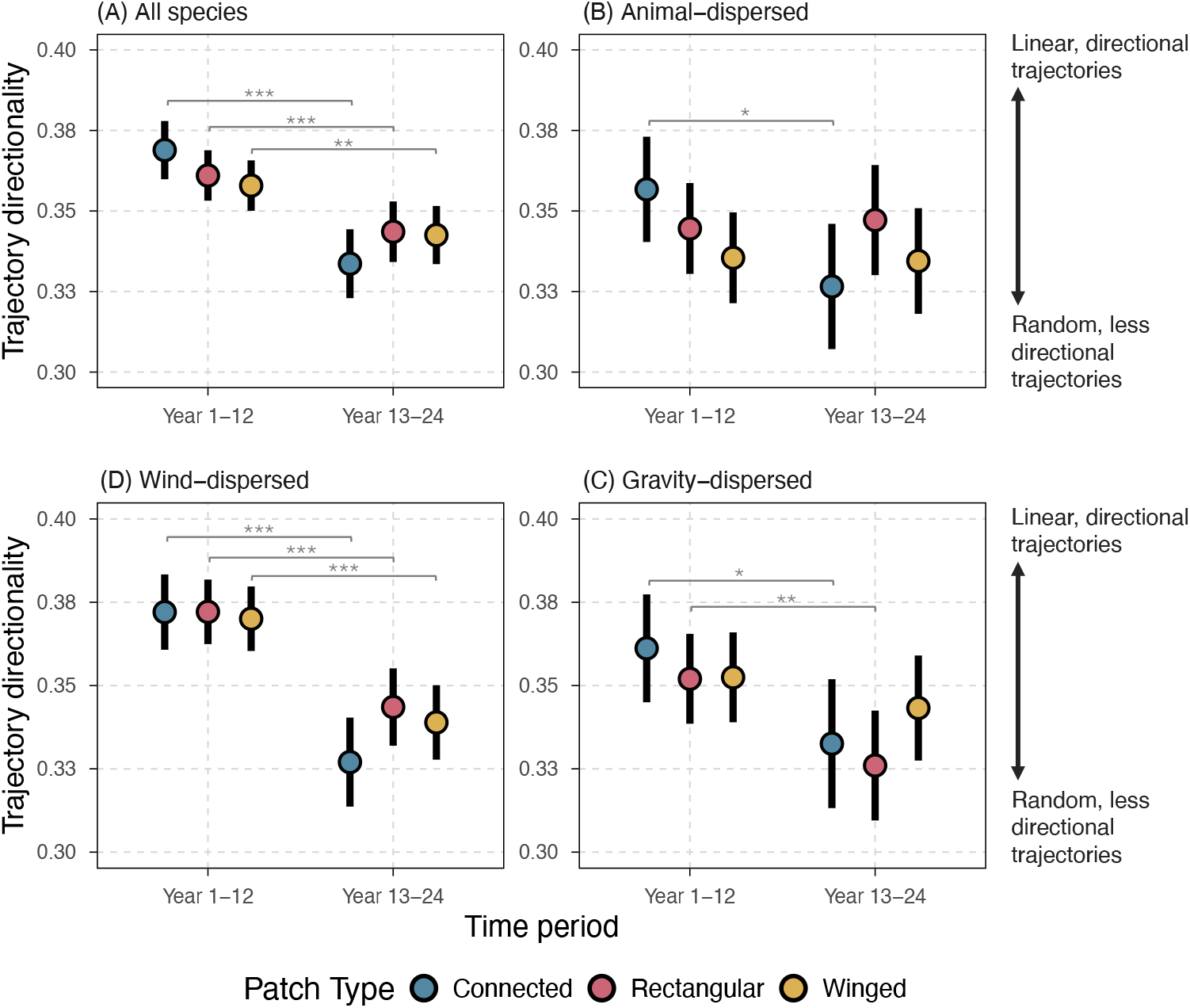
Directionality in assembly within a patch: Community trajectory directionality of (A) the entire plant community, (B) animal-dispersed species, (C) wind-dispersed species, and (D) gravity-dispersed species in the first half versus the second half of the time series. Higher values indicate more linear, directional changes in community composition, while smaller values indicate more random changes in composition. Solid points and error bars represent model predictions and 95% CI.

Patch connectivity and edge amount also did not affect directionality for any subset of dispersal modes (Table S7, S8). Across all patch types, however, the overall effect of time differed among dispersal modes (Table S9). For communities of animal-dispersed species, trajectories were similar in directionality in both earlier and later parts of the time series, except for connected patches where trajectories were less directional in the later part of the time series (Figure 3B; Table S9). For communities of gravity-dispersed species, trajectories in winged patches did not differ in directionality between time periods, but trajectories in connected and rectangular patches were less directional in the second half of the time series compared to the first half (Figure 3D; Table S12). Communities of wind-dispersed species in all patch types were more directional in the first half of the time series and less directional in the second half (Figure 3C; Table S12).

### Divergence in Composition Between Patches

All patch types diverged in composition across time (i.e., increase in spatial beta diversity over time), with a 10% increase in dissimilarity between patches over two decades (95% CI: 6.9%, 11.7% increase in dissimilarity from years 1 to 21; Figure 4A; Table S10-S12). Across time, patches with high amounts of edge habitat (connected and winged) were closer in composition to each other than patches differing in edge amount. Connected patches were 7% more compositionally similar to winged patches than rectangular patches, which have low amounts of edge habitat (95% CI: 4.1%, 10.0%), and winged patches were 4.3% more compositionally similar to connected patches than rectangular patches (95% CI: 1.6%, 6.8%) (Figure 4A; Table S11, S12).

**Figure 4.**
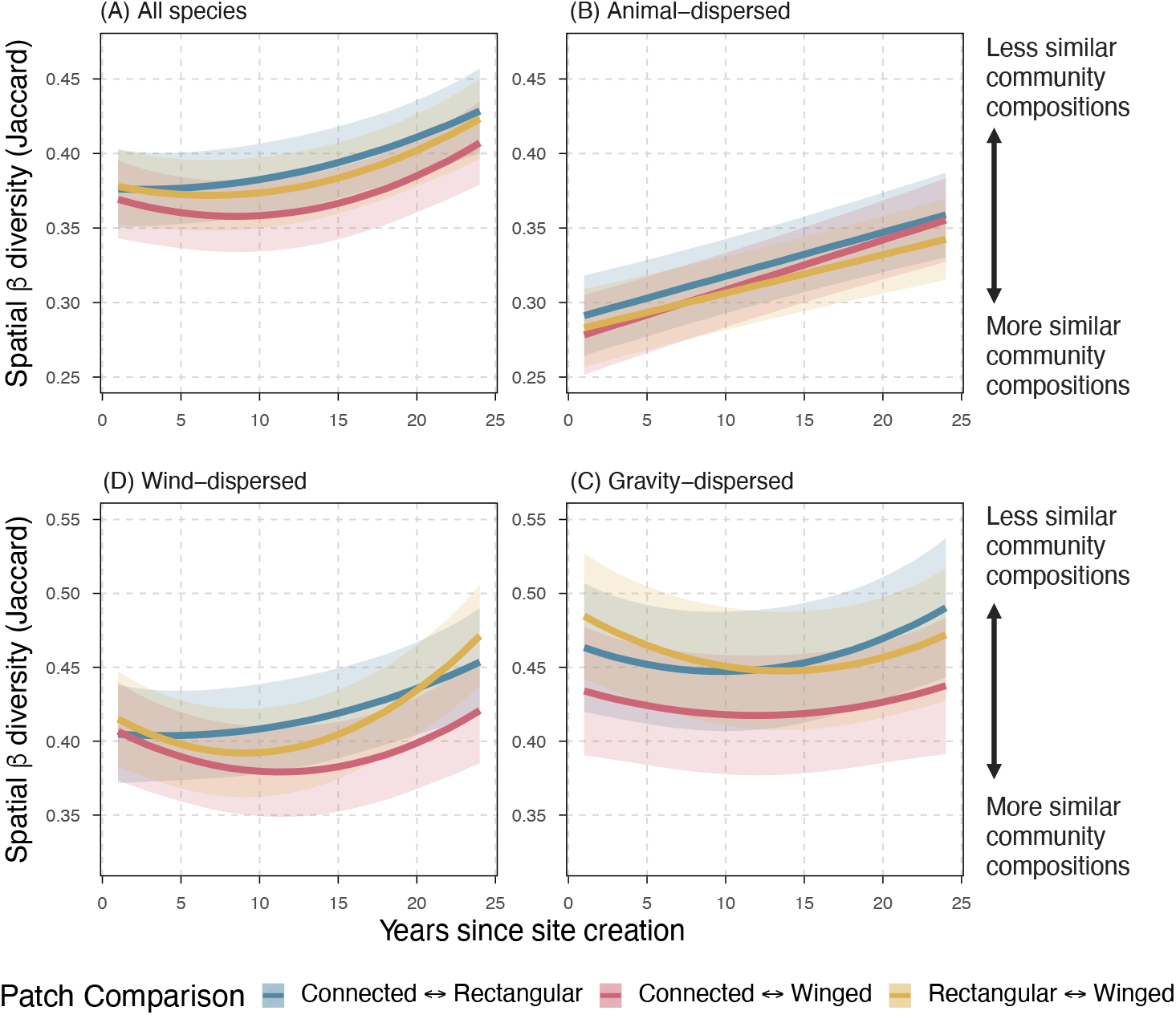
Divergence in composition between patches: Spatial beta diversity between pairs of patch types of (A) the entire plant community, (B) animal-dispersed species, (C) wind-dispersed species, and (D) gravity-dispersed species. The response variable is the Jaccard dissimilarity between two patches within a block during a single year. Higher values indicate less similar community compositions between patch pairs and lower values indicate more similar community compositions between patch pairs. Solid lines represent model predictions and shaded areas represent 95% CI. Note that y-axis scales differ among panels.

The rate of community divergence across time and comparisons between patch types differed among dispersal modes. For communities of animal-dispersed species, rectangular patches were more similar in composition to winged patches than connected patches, but this effect was small and there were no other pairwise differences between patch types. Across time, communities of animal-dispersed species diverged between all patch types, but at similar rates (Figure 4B; Table S11, S12).

For communities of wind-dispersed species, all patch types diverged in composition over time, and both connectivity and edge amount contributed to dissimilarity. At the midpoint of the time series, communities of wind-dispersed species were most dissimilar in patches that differed in both connectivity and edge amount (connected and rectangular patches). However, by the end of the time series, connectivity no longer contributed significantly to dissimilarity in wind-dispersed species composition: patches with similar edge amount were the most similar in composition (Figure 4C; Table S11, S12).

In contrast, communities of gravity-dispersed species neither converged nor diverged between most patch types, except for connected and rectangular patches which slightly diverged in composition over time. Patches with similar edge amounts (connected and winged) tended to be more similar in composition of gravity-dispersed species than patches that differed in edge amount (Figure 4D; Table S11, S12).

## Discussion

We demonstrate that spatial and temporal aspects of plant community assembly depend on landscape connectivity and edge amount over long time frames. Communities in connected patches experienced less temporal variability in composition compared to communities in unconnected patches, driven by a higher number of species persisting across years in connected patches. Although connectivity was important for temporal compositional changes within patches, spatial turnover between patches and directionality in assembly were more strongly impacted by the amount of edge habitat in a patch and duration of assembly. Our work experimentally isolates the role of connectivity in community assembly, avoiding spurious conclusions that could arise if connectivity were conflated with edge effects. By experimentally overcoming challenges that have confounded past studies, we clarify longstanding theoretical questions about assembly responses to connectivity-mediated dispersal, showing that landscape connectivity structures compositional dynamics in plant communities.

Plant communities in connected patches experienced less interannual change in community composition compared to unconnected patches (Figure 2). While theory predicts that intermediate levels of connectivity can stabilize communities (18), previous empirical research has found mixed results (41–44), which may be due in part to confounding connectivity with habitat or edge amount, conducting short-term studies, or differences in the composition of the habitat matrix between patches or the spatial scale of species dispersal ability relative to the landscape. Our results, which disentangle connectivity from other landscape features over a long time period, demonstrate that connectivity decreases temporal variability in local community composition, which may be due to spatial insurance from connectivity increasing species persistence in a patch. When partitioning interannual change into the number of species being gained and lost and the number of species staying present between consecutive years, we found that connected patches had a higher number of species that persisted from year to year compared to unconnected patches (Figure S2). Not only are species more likely to colonize connected patches earlier (40), but, as we show here, they are also likely to persist across years, reducing fluctuations in community composition in connected patches. By increasing the likelihood of dispersal between patches, connectivity can increase spatial insurance (45), which in our study may have conferred a greater compositional stability over time (18, 46).

Importantly, our experiment controlled for edge amount in a patch, unlike natural landscapes where edge habitat amount covaries with connectivity (47). We found that edge amount drove compositional divergence between patch types more than connectivity, suggesting that research that does not control for this relationship may misattribute these edge effects on assembly as connectivity effects. In our study, patch types with the same amount of edge habitat (connected and winged patches) were the most similar in composition over time, whereas patch types differing in both edge amount and connectivity (connected and rectangular) were the most dissimilar in composition (Figure 4A). While connectivity may have contributed in part to the compositional dissimilarity between connected and rectangular patches, patches with the same amount of edge habitat but differing in connectivity (connected and winged) were most similar in composition to each other, indicating that edge amount in a patch contributed more strongly to composition than connectivity. Because the patches in our experiment are open savanna habitat with plantation pine forest as the matrix between patches, areas near edges within patches tend to be shadier with higher amounts of leaf litter due to the dense pine overstory (48, 49). These distinct abiotic conditions at habitat edges may have led to compositional changes in patches with high amounts of edge through altered patterns in seed dispersal (50), establishment (49), and reproduction (49, 51).

The strong changes in assembly dynamics across time demonstrate the value of long-term research, as interpretations from our data based on shorter time frames would be insufficient to predict long-term community assembly. At the beginning of assembly, communities experienced high rates of interannual change in composition as species arrived via dispersal from other communities or emerged from the soil seed bank (29, 30). The high rate of compositional change early in assembly also tended to be more directional, indicating that deterministic processes may have played an important role in early dynamics. Over time, the rate of compositional change decreased in communities as more species arrived from the regional species pool and the direction of trajectory change became more random, suggesting that community change later in assembly may have resulted more from stochastic fluctuations in species occurrences (31). This trend may have emerged due to common, well-dispersing species arriving in habitat patches relatively early in the assembly process, while uncommon or rare species may arrive later and/or be more likely to go extinct, resulting in a pattern where compositional change shifts from more directional to more stochastic over longer timespans of community assembly.

Our work aligns with past research, both in our system and elsewhere, that shows that dispersal-limited species exhibit stronger responses to connectivity that persist over time (15, 35). Communities of wind and gravity-dispersed species tended to have stronger responses to connectivity and edge amount compared to animal-dispersed communities, with larger differences in spatial beta diversity and interannual compositional change between patch types (Figure 2, 3). Although vertebrates can be affected by connectivity at the scale of our experiment (39), they may have more frequent long-distance dispersal events compared to other groups, making animal-dispersed species less restricted in movement between patches (36). In contrast, connectivity may have especially strong effects on increasing the dispersal of comparatively poorer dispersers such as wind and gravity-dispersed species (37). Wind and gravity-dispersed communities also showed more distinct changes across time in assembly, with interannual compositional change decreasing for both abiotic dispersal mode communities and directionality of changes becoming more random for wind-dispersed communities later in assembly. These differences in responses between species with difference seed dispersal modes demonstrate the importance of using functional traits related to movement ability for explaining variation in responses to connectivity.

## Conclusion

Understanding how connectivity loss influences community assembly clarifies how regional dynamics affect community assembly and is significant for global conservation (8, 52). We experimentally disentangle connectivity from other landscape alterations, overcoming the challenges of studying long-term regional influences in local assembly. Our findings show that landscape connectivity and habitat edges regulate local community assembly, specifically for less-studied aspects of assembly like temporal variability in composition. Connectivity decreased temporal variability in composition, suggesting that spatial insurance that can buffer communities against environmental perturbations. Importantly, these trends may have been missed in studies of shorter time periods or without controlling for confounding variables such as edge amount, as is characteristic of prior work in this area (9, 30, 41, 53). Our work has widespread implications for biodiversity conservation and restoration. Anthropogenic land-use change leading to increased edge effects and a loss of habitat connectivity results in spatial and temporal changes in local community assembly.

## Materials and Methods

### Experimental Design

We conducted this study within a long-term landscape corridor experiment at the Savannah River Site (SRS, 33°21’46.5”N 81°40’58.7”W), a National Environmental Research Park in Aiken and Barnwell counties, South Carolina, USA. This experiment contains replicate landscapes (‘blocks’) created to test the effects of connectivity independent of habitat amount and edge amount. Each block was initiated by clearing mature pine plantation forest to create open habitat patches that are being restored to longleaf pine savanna. Eight blocks were created in winter 2000 and two additional blocks were created in winter 2007 (n = 10 blocks total); however, two of the original blocks were decommissioned in winter 2007 and one block was destroyed in 2015 by a windstorm. Therefore, six blocks were active for the entire time series (24 years), while the other four were active for shorter time periods (7-17 years). This temporal staggering of experimental landscape initiation creates a stronger study design for separating the effects of duration from those due to particular temporal events (e.g., climate).

Each block contains a 1-ha center patch surrounded by four 1.375 ha peripheral patches that differ in connectivity to the center patch and in the amount of edge habitat (Figure 1A). One peripheral patch is connected to the center patch by a 150 m × 25 m corridor (‘connected patch’). The other three peripheral patches are unconnected from the center patch and are either a 100 m × 137.5 m rectangle (‘rectangular patch’) or 100 m × 100 m with two 75 m × 25 m wings extending from opposite sides of the patch (‘winged patch’). Blocks are randomly assigned to contain either a duplicate winged patch or a duplicate rectangular patch. Winged patches have a similar edge amount as connected patches but differ in connectivity to the center patch, testing for connectivity effects. Rectangular patches and winged patches are both isolated from the center patch but differ in edge amount, testing for edge effects.

The habitat patches were created by clearing pine plantation forest, creating a distinct contrast between open habitat and forested matrix and providing the ability to track community assembly from the initiation of habitat restoration (Figure 1B). Habitat patches are managed with periodic low intensity prescribed fire and removal of woody encroachment, which aligns with historic fire regimes and management of longleaf pine savanna (54). This management allows patches to progress towards longleaf pine savanna communities, which is the historic habitat type for this region.

### Data collection

Plant species occurrence was surveyed annually each summer from 2001-2024 in all connected, winged, and rectangular patches, except for 2004 for all blocks, 2005-2006 for two of the eight active blocks, and 2013 where the duplicate patch type was not sampled in four of the eight active blocks. The entire patch area was systematically surveyed, and all vascular plants were identified to the species level, except for 4% of observations that were grouped at the genus level. See (40) for additional sampling details.

Each observed species was categorized as an animal-dispersed, wind-dispersed, or gravity-dispersed species, which are the three primary classes of seed dispersal observed in our experiment (see (40) for detailed methods). Animal dispersal included dispersal by vertebrates (i.e., mammals, birds) and invertebrates (i.e., ants). Gravity dispersal included species without any structures for assisted dispersal and species with ballistic dispersal. Species were categorized by botanical experts (including EID, LAB, CRW, MAB) using the Kew Garden Seed Information Database (55), primary literature describing dispersal of species, and morphological field observations. For the 4% of observations that were grouped at the genus level, all possible species within each genus had the same primary dispersal mode, so the genus was able to be assigned to a single dispersal mode.

### Statistical Analysis

First, we used community trajectory analysis to analyze how communities within each patch were changing temporally. Community trajectory analysis is a method of calculating geometric properties of temporal changes in communities using a multivariate space (56). We used Jaccard dissimilarity in community composition among all individual patches across time to specify the multivariate space and visualized community trajectories using a principal coordinates analysis (PCoA) (*ape* package v.5.8-1) (Figure S2) (57). To measure temporal variability in community composition within each patch, we calculated the distance of community change between consecutive years of sampling within each patch using the *trajectoryLengths* function from the *ecotraj* package v.1.2.0 (56). Because our experimental blocks were created in two different years, we used year since site creation as a predictor in models rather than calendar year. To evaluate if our response variable, trajectory length between consecutive years, was linear or nonlinear throughout time, we fit two generalized linear mixed effect models (GLMMs), both a linear model and a model with time as a quadratic term. We used AICc model selection to rank the linear and quadradic GLMMs against a null model (model with no fixed effects). The quadratic model with patch type, time, and the interaction of patch type and time as fixed effects was the best fit, with block as a random intercept (Table S1).

Interannual trajectory distance within a patch could increase due to species gains and losses or decrease due to a large set of core species persisting from year to year, buffering compositional changes. We partitioned compositional change between consecutive surveys within each patch into these parts by calculating the total number of gains and losses in a community between consecutive surveys (i.e., 0 → 1 or 1 → 0) and the total number of species persisting between two surveys (i.e., 1 → 1). To model the total number of gains and losses and the total number of persisting species, we used a single generalized additive model (GAMs) per response variable.

We chose to use GAMs because our initial attempts to fit GLMMS exhibited poor fit and convergence issues. Each GAM included patch type as a predictor and patch type specific smooth of time (k = 5) to capture patterns across time.

Next, we used a second community trajectory analysis metric to quantify whether communities within patches are changing linearly or randomly throughout time. We measured directionality using the *trajectoryDirectionality* function from the *ecotraj* package, which is a metric that uses the angles between segments of a trajectory to measure whether a community is changing linearly or randomly in multivariate space (56). Values of 1 indicate a completely straight trajectory, while values closer to 0 indicate a trajectory with sharp changes in direction. Because this metric required three or more years to be calculated and we expected directionality to change throughout assembly, we chose to calculate directionality for each community across two time periods, once across the first 12 years of assembly and then for the second 12 years of assembly. We fit one GLMM with trajectory directionality as the response variable and the following predictor variables: time period, patch type, and the interaction between time period and patch type as fixed effects and block as a random intercept.

We assessed convergence or divergence patterns between patch types by calculating the dissimilarity in community composition over time. For each year of sampling, we calculated the Jaccard dissimilarity between each pair of patches (connected vs. winged, connected vs. rectangular, winged vs. rectangular) within each block using the *vegdist* function from the *vegan* package v.2.7-2 (58). Decreasing values over time indicate convergence in composition, while increasing values over time indicate divergence in composition. We used AICc model selection to rank a linear, quadratic, and null model, and the quadratic model was the best fit (Table S10).

Lastly, we repeated each analysis for each dispersal mode. We divided the community into animal-dispersed species, wind-dispersed species, and gravity-dispersed species and repeated calculations of interannual trajectory segment length, trajectory directionality, and convergence and divergence patterns with each subset of species. Because our goal was to evaluate trends within a dispersal mode, and due to model limitations, we fit models individually for each dispersal mode, using the same methods as above to determine the best fit model for each response variable (AICc comparisons in Table S1, S10).

All analyses were conducted in R v.4.5.2 (59). We fit generalized linear models using the glmmTMB package v.1.1.13 (60, 61), checked for model fit using the DHARMa package v.0.4.7 (62) and performance package v.0.15.2 (63), conducted Tukey posthoc tests using the emmeans package v.2.0.0 (64), and plotted using the ggeffects package v.2.3.1 (65) and ggplot v.4.0.0 (66).

## Acknowledgments

We thank John Blake, Ed Olson, Andy Horcher, Jim Segar, James Garabedian, Sabrie Breland, Benjamin Overlie, and the fire management crew of the USDA Forest Service for their role in implementing and maintaining the experimental habitat corridors. We also thank Doug Levey, John Orrock, and Julian Resasco for their leadership on the experiment and many undergraduate students and technicians for assistance over the years. Funding was provided by the National Science Foundation (Awards DEB-1912729 and DEB-1913501) and the Department of Energy—Savannah River Operations Office through the U.S. Forest Service—Savannah River under Interagency Agreement DE-89303720SEM000037. KAH was funded by the National Science Foundation Graduate Research Fellowship Program (award number 2235783).

