## Supplemental Information for "Experimental landscape connectivity decreases temporal variability in communities over 24 years of assembly"

**Figures**

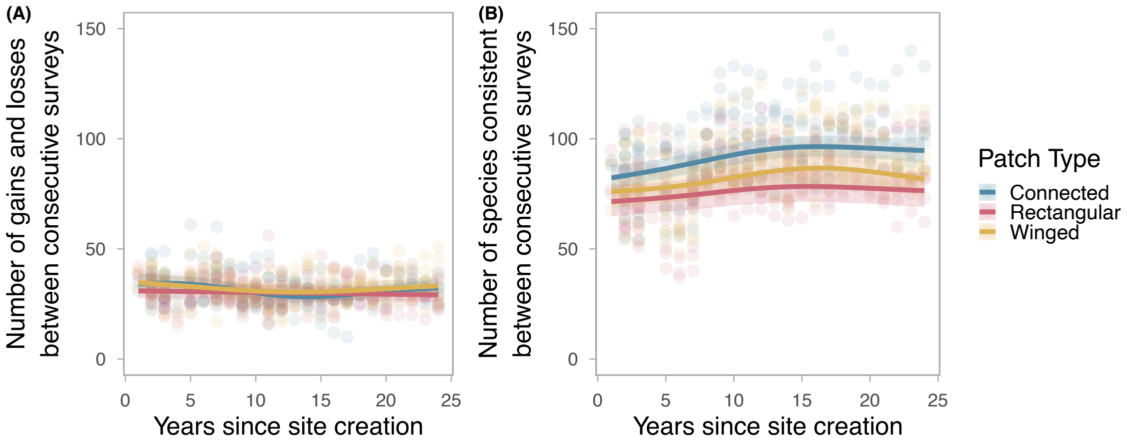


**Fig. S1.** (A) Number of total gains and losses of species in a patch between consecutive surveys (i.e., turnover between years). (B) Number of species persisting in a patch between consecutive surveys. Points represent values from a single patch and shaded regions are 95% CI.

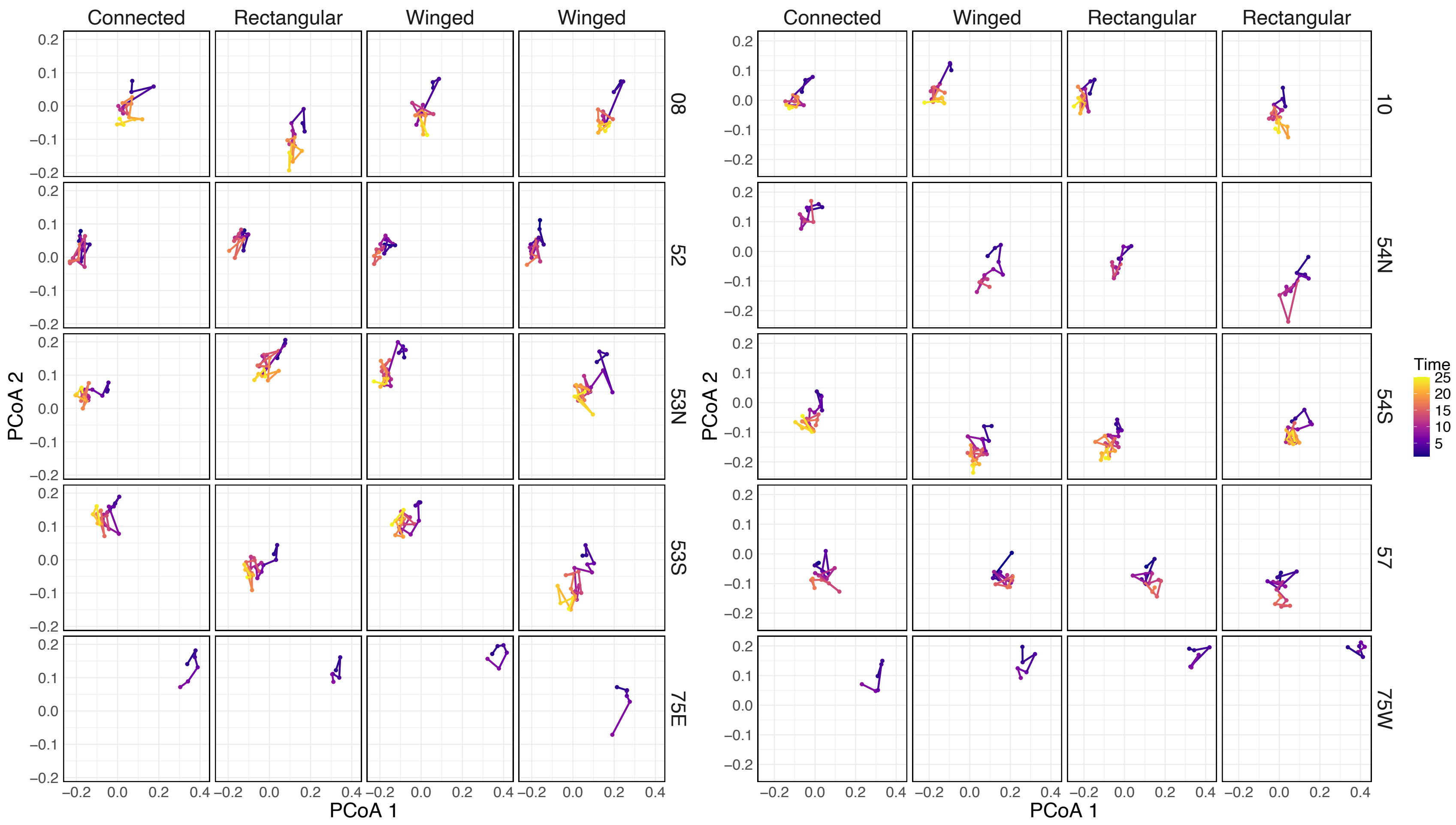


**Fig. S2.** PCoA of community trajectories in patches. Points represent PCoA scores for the composition of a patch in one year and lines connect consecutive years. Each row is labelled with the experimental block that the patch is located in

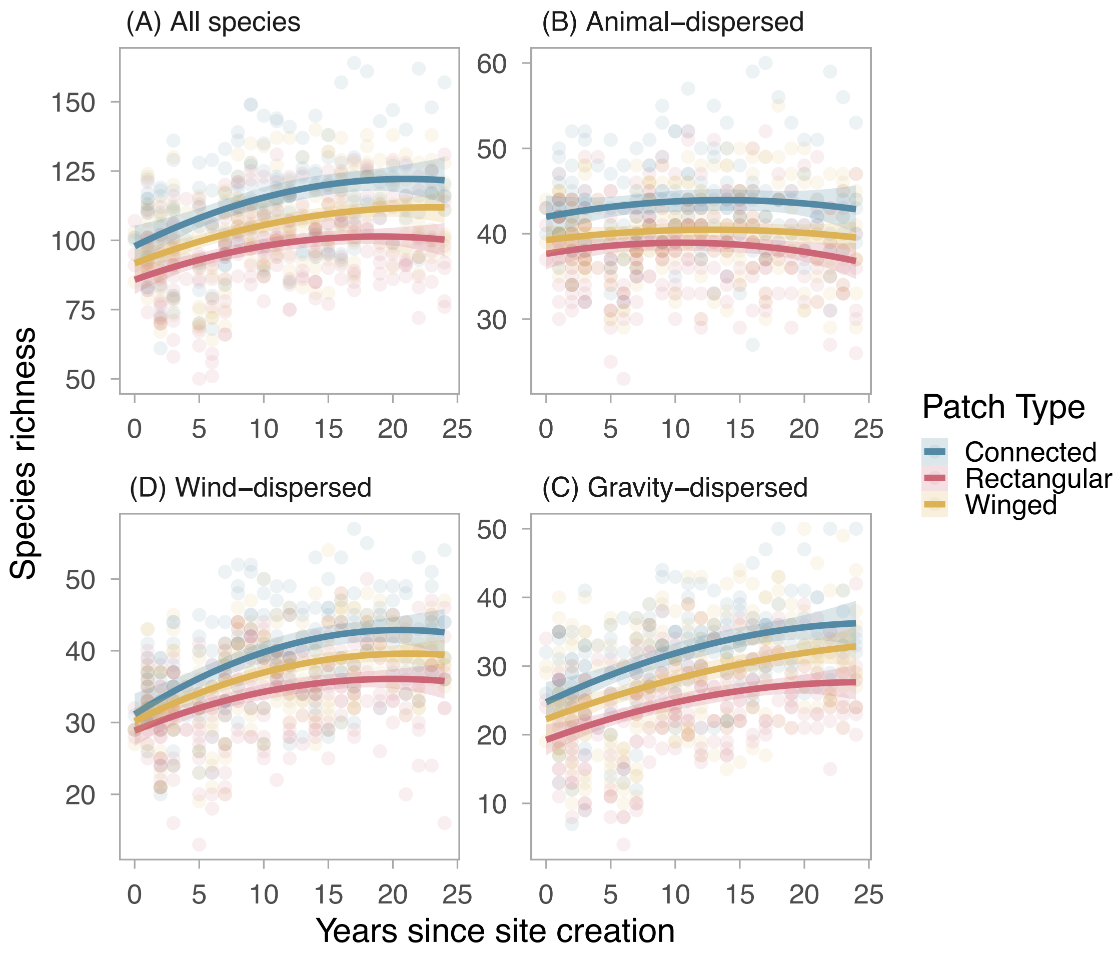


**Fig. S3.** Species richness across time for A) the entire plant community, B) animal-dispersed species, C) wind-dispersed species, and D) gravity-dispersed species. Points represent the species richness of one patch during one sampling point, lines represent model predictions (years since site creation as a quadratic term), and shaded regions represent 95% CI. Note that axis scales differ between plots.

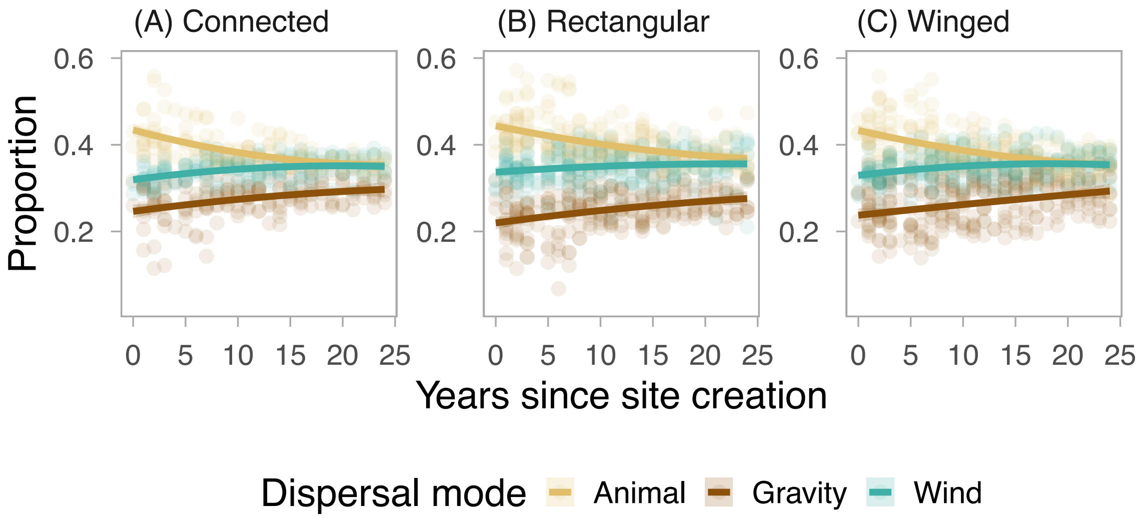


**Fig. S4.** The proportion of animal, gravity, and wind-dispersed species over time in A) connected patches, B) rectangular patches, and C) winged patches. Each point represents the proportion of species in one dispersal mode group to the total number of plant species in a patch at one time point.

### Tables

**Table S1.** AICc model comparison for the effect of patch type (connected, winged, rectangular) and time on community trajectory segment length between consecutive surveys (interannual community change). K is the number of parameters in the model and LL is the log likelihood. If a model was within 2 ΔAICc of the top ranked model, we interpreted results from the most parsimonious model.

| Dispersal Mode | Model | Model Formula | K | LL | AICc | Delta AICc | Cumulative Weight |
|---|---|---|---|---|---|---|---|
| All Species | Quadratic | $\sim \beta_0 + \beta_1$(Patch) + $\beta_2$(Time) + $\beta_3$(Patch*Time) + $\beta_4$(Time$^2$) + $\beta_5$(Patch*Time$^2$) + $b_{\text{Block}}$ + $b_{\text{PatchReplicate(Block)}}$ | 12 | 959.398 | -1894.288 | 0.000 | 1.000 |
| | Linear | $\sim \beta_0 + \beta_1$(Patch) + $\beta_2$(Time) + $\beta_3$(Patch*Time) + $b_{\text{Block}}$ + $b_{\text{PatchReplicate(Block)}}$ | 9 | 945.661 | -1873.030 | 21.258 | 1.000 |
| | Null | $\sim \beta_0$ + $b_{\text{Block}}$ + $b_{\text{PatchReplicate(Block)}}$ | 4 | 928.889 | -1849.713 | 44.575 | 1.000 |
| Animal-Dispersed | Null | $\sim \beta_0$ + $b_{\text{Block}}$ + $b_{\text{PatchReplicate(Block)}}$ | 4 | 808.136 | -1608.207 | 0.000 | 0.758 |
| | Linear | $\sim \beta_0 + \beta_1$(Patch) + $\beta_2$(Time) + $\beta_3$(Patch*Time) + $b_{\text{Block}}$ + $b_{\text{PatchReplicate(Block)}}$ | 9 | 811.917 | -1605.543 | 2.664 | 0.958 |
| | Quadratic | $\sim \beta_0 + \beta_1$(Patch) + $\beta_2$(Time) + $\beta_3$(Patch*Time) + $\beta_4$(Time$^2$) + $\beta_5$(Patch*Time$^2$) + $b_{\text{Block}}$ + $b_{\text{PatchReplicate(Block)}}$ | 12 | 813.459 | -1602.410 | 5.797 | 1.000 |
| Gravity-Dispersed | Quadratic | $\sim \beta_0 + \beta_1$(Patch) + $\beta_2$(Time) + $\beta_3$(Patch*Time) + $\beta_4$(Time$^2$) + $\beta_5$(Patch*Time$^2$) + $b_{\text{Block}}$ + $b_{\text{PatchReplicate(Block)}}$ | 12 | 624.118 | -1223.729 | 0.000 | 0.992 |
| | Linear | $\sim \beta_0 + \beta_1$(Patch) + $\beta_2$(Time) + $\beta_3$(Patch*Time) + $b_{\text{Block}}$ + $b_{\text{PatchReplicate(Block)}}$ | 9 | 616.148 | -1214.005 | 9.725 | 1.000 |
| | Null | $\sim \beta_0$ + $b_{\text{Block}}$ + $b_{\text{PatchReplicate(Block)}}$ | 4 | 596.200 | -1184.335 | 39.395 | 1.000 |
| Wind-Dispersed | Quadratic | $\sim \beta_0 + \beta_1$(Patch) + $\beta_2$(Time) + $\beta_3$(Patch*Time) + $\beta_4$(Time$^2$) + $\beta_5$(Patch*Time$^2$) + $b_{\text{Block}}$ + $b_{\text{PatchReplicate(Block)}}$ | 12 | 729.863 | -1435.218 | 0.000 | 1.000 |
| | Linear | $\sim \beta_0 + \beta_1$(Patch) + $\beta_2$(Time) + $\beta_3$(Patch*Time) + $b_{\text{Block}}$ + $b_{\text{PatchReplicate(Block)}}$ | 9 | 710.771 | -1403.252 | 31.967 | 1.000 |
| | Null | $\sim \beta_0$ + $b_{\text{Block}}$ + $b_{\text{PatchReplicate(Block)}}$ | 4 | 695.885 | -1383.707 | 51.512 | 1.000 |

**Table S2.** Anova type III results of the GLMM testing the effect of patch type (connected, winged, rectangular) and time on community trajectory segment length between consecutive surveys (interannual community change). Bolded terms indicate significant predictors ($p$ < 0.05).

| Dispersal mode | Top Model | Variable | Chisq | df | p.value |
|---|---|---|---|---|---|
| All Species | Quadratic | **Intercept** | **314.556** | **1** | **0.000** |
| | | **Patch Type** | **12.658** | **2** | **0.002** |
| | | **Time** | **15.499** | **1** | **0.000** |
| | | **Time^2** | **11.745** | **1** | **0.001** |
| | | Patch Type:Time | 1.777 | 2 | 0.411 |
| | | Patch Type:Time^2 | 3.598 | 2 | 0.165 |
| Animal-Dispersed | Null | **Intercept** | **558.086** | **1** | **0.000** |
| Gravity-Dispersed | Quadratic | **Intercept** | **106.891** | **1** | **0.000** |
| | | Patch Type | 5.203 | 2 | 0.074 |
| | | **Time** | **25.319** | **1** | **0.000** |
| | | **Time^2** | **4.789** | **1** | **0.029** |
| | | Patch Type:Time | 4.450 | 2 | 0.108 |
| | | Patch Type:Time^2 | 1.232 | 2 | 0.540 |
| Wind-Dispersed | Quadratic | **Intercept** | **256.692** | **1** | **0.000** |
| | | **Patch Type** | **7.550** | **2** | **0.023** |
| | | **Time** | **16.640** | **1** | **0.000** |
| | | **Time^2** | **14.650** | **1** | **0.000** |
| | | Patch Type:Time | 3.452 | 2 | 0.178 |
| | | Patch Type:Time^2 | 1.909 | 2 | 0.385 |

**Table S3.** Emmeans pairwise comparisons of the effects of patch type on community trajectory segment length across time. Contrasts are at the midpoint of the time series. Bolded terms indicate significant predictors ($p < 0.05$).

| Dispersal mode | contrast | estimate | SE | df | z.ratio | p.value |
|---|---|---|---|---|---|---|
| All Species | **Connected - Rectangular** | **-0.039** | **0.011** | **Inf** | **-3.536** | **0.001** |
| | **Connected - Winged** | **-0.026** | **0.011** | **Inf** | **-2.373** | **0.046** |
| | Rectangular - Winged | 0.013 | 0.010 | Inf | 1.296 | 0.397 |
| Gravity-Dispersed | Connected - Rectangular | -0.047 | 0.021 | Inf | -2.279 | 0.059 |
| | Connected - Winged | -0.025 | 0.020 | Inf | -1.244 | 0.427 |
| | Rectangular - Winged | 0.021 | 0.019 | Inf | 1.141 | 0.489 |
| Wind-Dispersed | **Connected - Rectangular** | **-0.044** | **0.016** | **Inf** | **-2.742** | **0.017** |
| | Connected - Winged | -0.023 | 0.016 | Inf | -1.451 | 0.315 |
| | Rectangular - Winged | 0.021 | 0.015 | Inf | 1.422 | 0.329 |

**Table S4.** Results of the generalized additive model (GAM) for the effect of patch type and time on the total number of species gains and losses between years. Bolded terms indicate significant predictors (*p* < 0.05).

| Variable | estimate | SE | z value | p.value |
|---|---|---|---|---|
| **(Intercept)** | **3.437** | **0.036** | **96.213** | **0.000** |
| patch_typeRectangular | -0.033 | 0.043 | -0.747 | 0.455 |
| patch_typeWinged | 0.024 | 0.043 | 0.554 | 0.580 |
| **Variable** | **edf** | **Ref.df** | **Chi.sq** | **p.value** |
| **Time:Connected Patch** | **3.197** | **3.651** | **19.321** | **0.000** |
| Time:Rectangular Patch | 1.001 | 1.002 | 1.493 | 0.222 |
| **Time:Winged Patch** | **2.525** | **3.030** | **10.982** | **0.012** |

**Table S5.** Results of the generalized additive model (GAM) for the effect of patch type and time on the total number of persisting species between years. Bolded terms indicate significant predictors ($p < 0.05$).

| Variable | estimate | SE | z value | p.value |
|---|---|---|---|---|
| **(Intercept)** | **4.520** | **0.067** | **67.842** | **0.000** |
| **patch_typeRectangular** | **-0.190** | **0.042** | **-4.540** | **0.000** |
| **patch_typeWinged** | **-0.112** | **0.042** | **-2.676** | **0.007** |
| Variable | edf | Ref.df | Chi.sq | p.value |
| **Time:Connected Patch** | **2.596** | **3.107** | **33.448** | **0.000** |
| **Time:Rectangular Patch** | **2.458** | **2.962** | **15.075** | **0.002** |
| **Time:Winged Patch** | **3.007** | **3.495** | **36.246** | **0.000** |

**Table S6.** Emmeans pairwise comparisons of the effects of patch type on 1) the total number of species gains and losses between years and 2) total number of persisting species between years. Contrasts are at the midpoint of the time series. Bolded terms indicate significant predictors ($p < 0.05$).

| Response | contrast | estimate | SE | df | t.ratio | p.value |
|---|---|---|---|---|---|---|
| Number of gains and losses | Connected - Rectangular | -0.044 | 0.049 | 590.095 | -0.904 | 0.638 |
| | Connected - Winged | -0.057 | 0.051 | 590.095 | -1.106 | 0.511 |
| | Rectangular - Winged | -0.012 | 0.042 | 590.095 | -0.293 | 0.954 |
| Number of persisting species | **Connected - Rectangular** | **0.199** | **0.044** | **583.120** | **4.539** | **0.000** |
| | **Connected - Winged** | **0.115** | **0.044** | **583.120** | **2.616** | **0.025** |
| | Rectangular - Winged | -0.084 | 0.041 | 583.120 | -2.057 | 0.100 |

**Table S7.** Anova type III results of the GLMM testing the effect of patch type (connected, winged, rectangular) and time period (Year 1-12 of time series, Year 13-24 of time series) on community trajectory directionality. Bolded terms indicate significant predictors ($p < 0.05$).

| Dispersal mode | Variable | Chisq | df | p.value |
|---|---|---|---|---|
| All Species | Patch Type | 4.446 | 2 | 0.108 |
| | **Time Period** | **29.767** | **1** | **0.000** |
| | **Patch Type:Time Period** | **6.694** | **2** | **0.035** |
| Animal-Dispersed | Patch Type | 4.875 | 2 | 0.087 |
| | **Time Period** | **6.520** | **1** | **0.011** |
| | Patch Type:Time Period | 5.412 | 2 | 0.067 |
| Gravity-Dispersed | Patch Type | 0.943 | 2 | 0.624 |
| | **Time Period** | **5.231** | **1** | **0.022** |
| | Patch Type:Time Period | 1.977 | 2 | 0.372 |
| Wind-Dispersed | Patch Type | 0.136 | 2 | 0.934 |
| | **Time Period** | **29.633** | **1** | **0.000** |
| | Patch Type:Time Period | 2.637 | 2 | 0.268 |

**Table S8.** Emmeans pairwise comparisons of the effects of patch type on community trajectory directionality within a time period. Bolded terms indicate significant predictors ($p < 0.05$).

| Dispersal mode | Time | contrast | estimate | SE | df | z.ratio | p.value |
|---|---|---|---|---|---|---|---|
| All Species | Year 1-12 | Connected - Rectangular | 0.008 | 0.005 | Inf | 1.489 | 0.296 |
| | | Connected - Winged | 0.011 | 0.005 | Inf | 2.083 | 0.093 |
| | | Rectangular - Winged | 0.003 | 0.005 | Inf | 0.654 | 0.790 |
| | Year 13-24 | Connected - Rectangular | -0.010 | 0.006 | Inf | -1.556 | 0.265 |
| | | Connected - Winged | -0.009 | 0.006 | Inf | -1.421 | 0.330 |
| | | Rectangular - Winged | 0.001 | 0.006 | Inf | 0.181 | 0.982 |
| Animal-Dispersed | Year 1-12 | Connected - Rectangular | 0.012 | 0.010 | Inf | 1.262 | 0.417 |
| | | Connected - Winged | 0.021 | 0.010 | Inf | 2.207 | 0.070 |
| | | Rectangular - Winged | 0.009 | 0.009 | Inf | 1.043 | 0.550 |
| | Year 13-24 | Connected - Rectangular | -0.021 | 0.012 | Inf | -1.774 | 0.178 |
| | | Connected - Winged | -0.008 | 0.011 | Inf | -0.693 | 0.768 |
| | | Rectangular - Winged | 0.013 | 0.010 | Inf | 1.220 | 0.441 |
| Gravity-Dispersed | Year 1-12 | Connected - Rectangular | 0.009 | 0.010 | Inf | 0.887 | 0.648 |
| | | Connected - Winged | 0.009 | 0.010 | Inf | 0.844 | 0.676 |
| | | Rectangular - Winged | 0.000 | 0.009 | Inf | -0.048 | 0.999 |
| | Year 13-24 | Connected - Rectangular | 0.007 | 0.012 | Inf | 0.527 | 0.858 |
| | | Connected - Winged | -0.011 | 0.012 | Inf | -0.875 | 0.656 |
| | | Rectangular - Winged | -0.017 | 0.011 | Inf | -1.554 | 0.266 |
| Wind-Dispersed | Year 1-12 | Connected - Rectangular | 0.000 | 0.007 | Inf | -0.011 | 1.000 |
| | | Connected - Winged | 0.002 | 0.007 | Inf | 0.292 | 0.954 |
| | | Rectangular - Winged | 0.002 | 0.006 | Inf | 0.334 | 0.940 |
| | Year 13-24 | Connected - Rectangular | -0.017 | 0.008 | Inf | -2.013 | 0.109 |
| | | Connected - Winged | -0.012 | 0.008 | Inf | -1.477 | 0.302 |
| | | Rectangular - Winged | 0.005 | 0.007 | Inf | 0.630 | 0.804 |

**Table S9.** Emmeans pairwise comparisons of the effects of time period on community trajectory directionality within a patch type. Bolded terms indicate significant predictors ($p < 0.05$).

| Dispersal mode | Patch Type | contrast | estimate | SE | df | z.ratio | p.value |
|---|---|---|---|---|---|---|---|
| All Species | **Connected** | **(Year 1-12) - (Year 13-24)** | **0.035** | **0.006** | **Inf** | **5.456** | **0.000** |
| | **Rectangular** | **(Year 1-12) - (Year 13-24)** | **0.017** | **0.005** | **Inf** | **3.217** | **0.001** |
| | **Winged** | **(Year 1-12) - (Year 13-24)** | **0.015** | **0.005** | **Inf** | **2.933** | **0.003** |
| Animal-Dispersed | **Connected** | **(Year 1-12) - (Year 13-24)** | **0.030** | **0.012** | **Inf** | **2.554** | **0.011** |
| | Rectangular | (Year 1-12) - (Year 13-24) | -0.003 | 0.010 | Inf | -0.262 | 0.793 |
| | Winged | (Year 1-12) - (Year 13-24) | 0.001 | 0.010 | Inf | 0.105 | 0.917 |
| Gravity-Dispersed | **Connected** | **(Year 1-12) - (Year 13-24)** | **0.029** | **0.013** | **Inf** | **2.287** | **0.022** |
| | **Rectangular** | **(Year 1-12) - (Year 13-24)** | **0.026** | **0.010** | **Inf** | **2.499** | **0.012** |
| | Winged | (Year 1-12) - (Year 13-24) | 0.009 | 0.010 | Inf | 0.911 | 0.362 |
| Wind-Dispersed | **Connected** | **(Year 1-12) - (Year 13-24)** | **0.045** | **0.008** | **Inf** | **5.444** | **0.000** |
| | **Rectangular** | **(Year 1-12) - (Year 13-24)** | **0.029** | **0.007** | **Inf** | **4.131** | **0.000** |
| | **Winged** | **(Year 1-12) - (Year 13-24)** | **0.031** | **0.007** | **Inf** | **4.656** | **0.000** |

**Table S10.** AICc model comparison for the effect of patch type comparison (connected↔rectangular, connected↔winged, winged↔rectangular) and time on spatial β diversity (Jaccard dissimilarity). K is the number of parameters in the model and LL is the log likelihood. If a model was within 2 ΔAICc of the top ranked model, we interpreted results from the most parsimonious model.

| Dispersal Mode | Model | Model Formula | K | LL | AICc | Delta AICc | Cumulative Weight |
|---|---|---|---|---|---|---|---|
| All Species | Quadratic | $\sim \beta_0 + \beta_1(\text{PatchPair}) + \beta_2(\text{Time}) + \beta_3(\text{PatchPair*Time}) + \beta_4(\text{Time}^2) + \beta_5(\text{PatchPair*Time}^2) + b_{\text{Block}}$ | 11 | 2340.218 | -4658.275 | 0.000 | 1.000 |
| | Linear | $\sim \beta_0 + \beta_1(\text{PatchPair}) + \beta_2(\text{Time}) + \beta_3(\text{PatchPair*Time}) + b_{\text{Block}}$ | 8 | 2327.478 | -4638.869 | 19.406 | 1.000 |
| | Null | $\sim \beta_0 + b_{\text{Block}}$ | 3 | 2282.952 | -4559.889 | 98.386 | 1.000 |
| Animal-Dispersed | Quadratic | $\sim \beta_0 + \beta_1(\text{PatchPair}) + \beta_2(\text{Time}) + \beta_3(\text{PatchPair*Time}) + \beta_4(\text{Time}^2) + \beta_5(\text{PatchPair*Time}^2) + b_{\text{Block}}$ | 11 | 1929.623 | -3837.083 | 0.000 | 0.664 |
| | Linear | $\sim \beta_0 + \beta_1(\text{PatchPair}) + \beta_2(\text{Time}) + \beta_3(\text{PatchPair*Time}) + b_{\text{Block}}$ | 8 | 1925.905 | -3835.723 | 1.361 | 1.000 |
| | Null | $\sim \beta_0 + b_{\text{Block}}$ | 3 | 1875.296 | -3744.577 | 92.507 | 1.000 |
| Gravity-Dispersed | Quadratic | $\sim \beta_0 + \beta_1(\text{PatchPair}) + \beta_2(\text{Time}) + \beta_3(\text{PatchPair*Time}) + \beta_4(\text{Time}^2) + \beta_5(\text{PatchPair*Time}^2) + b_{\text{Block}}$ | 11 | 1577.467 | -3132.773 | 0.000 | 0.983 |
| | Linear | $\sim \beta_0 + \beta_1(\text{PatchPair}) + \beta_2(\text{Time}) + \beta_3(\text{PatchPair*Time}) + b_{\text{Block}}$ | 8 | 1570.383 | -3124.678 | 8.095 | 1.000 |
| | Null | $\sim \beta_0 + b_{\text{Block}}$ | 3 | 1544.664 | -3083.313 | 49.460 | 1.000 |
| Wind-Dispersed | Quadratic | $\sim \beta_0 + \beta_1(\text{PatchPair}) + \beta_2(\text{Time}) + \beta_3(\text{PatchPair*Time}) + \beta_4(\text{Time}^2) + \beta_5(\text{PatchPair*Time}^2) + b_{\text{Block}}$ | 11 | 1880.557 | -3738.952 | 0.000 | 1.000 |
| | Linear | $\sim \beta_0 + \beta_1(\text{PatchPair}) + \beta_2(\text{Time}) + \beta_3(\text{PatchPair*Time}) + b_{\text{Block}}$ | 8 | 1861.485 | -3706.882 | 32.070 | 1.000 |
| | Null | $\sim \beta_0 + b_{\text{Block}}$ | 3 | 1833.123 | -3660.230 | 78.722 | 1.000 |

**Table S11.** Anova type III results of the generalized linear mixed effects model (GLMM) testing the effect of patch type comparison (connected↔rectangular, connected↔winged, winged↔rectangular**)** and time on spatial β diversity (Jaccard dissimilarity). Bolded terms indicate significant predictors ($p < 0.05$).

| Dispersal mode | Top Model | Variable | Chisq | df | p.value |
|---|---|---|---|---|---|
| All Species | Quadratic | **Intercept** | **978.374** | **1** | **0.000** |
| | | **Patch Pair** | **22.853** | **2** | **0.000** |
| | | **Time** | **23.894** | **1** | **0.000** |
| | | Time^2 | 3.368 | 1 | 0.066 |
| | | Patch Pair:Time | 2.380 | 2 | 0.304 |
| | | Patch Pair:Time^2 | 1.249 | 2 | 0.536 |
| Animal-Dispersed | Linear | **Intercept** | **638.205** | **1** | **0.000** |
| | | **Patch Pair** | **7.088** | **2** | **0.029** |
| | | **Time** | **31.772** | **1** | **0.000** |
| | | Patch Pair:Time | 1.401 | 2 | 0.496 |
| Gravity-Dispersed | Quadratic | **Intercept** | **469.695** | **1** | **0.000** |
| | | **Patch Pair** | **18.388** | **2** | **0.000** |
| | | Time | 1.007 | 1 | 0.316 |
| | | **Time^2** | **4.777** | **1** | **0.029** |
| | | Patch Pair:Time | 4.599 | 2 | 0.100 |
| | | Patch Pair:Time^2 | 0.580 | 2 | 0.748 |
| Wind-Dispersed | Quadratic | **Intercept** | **697.937** | **1** | **0.000** |
| | | **Patch Pair** | **19.137** | **2** | **0.000** |
| | | **Time** | **11.215** | **1** | **0.001** |
| | | Time^2 | 2.338 | 1 | 0.126 |
| | | **Patch Pair:Time** | **7.663** | **2** | **0.022** |
| | | Patch Pair:Time^2 | 4.830 | 2 | 0.089 |

**Table S12.** Emmeans pairwise comparisons of the effects of patch pair comparisons on spatial β diversity (Jaccard dissimilarity) across time. Contrasts are at the midpoint of the time series. Bolded terms indicate significant predictors ($p$ < 0.05).

| Dispersal mode | contrast | estimate | SE | df | z.ratio | p.value |
|---|---|---|---|---|---|---|
| All Species | **(Connected-Rectangular) - (Connected-Winged)** | **0.025** | **0.005** | **Inf** | **4.715** | **0.000** |
| | (Connected-Rectangular) - (Rectangular-Winged) | 0.010 | 0.005 | Inf | 1.908 | 0.136 |
| | **(Connected-Winged) - (Rectangular-Winged)** | **-0.016** | **0.005** | **Inf** | **-3.188** | **0.004** |
| Animal-Dispersed | (Connected-Rectangular) - (Connected-Winged) | 0.009 | 0.005 | Inf | 1.781 | 0.176 |
| | **(Connected-Rectangular) - (Rectangular-Winged)** | **0.012** | **0.004** | **Inf** | **2.634** | **0.023** |
| | (Connected-Winged) - (Rectangular-Winged) | 0.003 | 0.004 | Inf | 0.708 | 0.759 |
| Gravity-Dispersed | **(Connected-Rectangular) - (Connected-Winged)** | **0.030** | **0.009** | **Inf** | **3.514** | **0.001** |
| | (Connected-Rectangular) - (Rectangular-Winged) | -0.001 | 0.008 | Inf | -0.146 | 0.988 |
| | **(Connected-Winged) - (Rectangular-Winged)** | **-0.031** | **0.008** | **Inf** | **-3.956** | **0.000** |
| Wind-Dispersed | **(Connected-Rectangular) - (Connected-Winged)** | **0.031** | **0.007** | **Inf** | **4.372** | **0.000** |
| | **(Connected-Rectangular) - (Rectangular-Winged)** | **0.017** | **0.007** | **Inf** | **2.511** | **0.032** |
| | (Connected-Winged) - (Rectangular-Winged) | -0.015 | 0.007 | Inf | -2.208 | 0.070 |
